# Clonazepam activates the transient receptor potential melastatin 8 (TRPM8) ion channel

**DOI:** 10.64898/2026.01.19.700423

**Authors:** Logan Elkin, Evgeny G. Chulkov, Jonathan Enders, Anthony D. Menzel, Sang-Kyu Park, Cheryl L. Stucky, Jonathan S. Marchant

## Abstract

Benzodiazepines represent a major class of drugs prescribed as treatments for anxiety, insomnia and seizures. They act by engaging GABA_A_ receptors in the central nervous system to depress neuronal excitability. Here, the polypharmacology of benzodiazepines was explored by studying their engagement of human TRP channels revealing clonazepam acts as a robust and selective activator of the TRPM8 ion channel. Clonazepam-evoked Ca^2+^ signals were observed in cells expressing human TRPM8 channels, and in mouse trigeminal neurons. These responses were completely blocked by pharmacological or genetic inhibition of TRPM8 function. This discovery likely explains why clonazepam is an effective treatment for the painful oral condition known as burning mouth disorder, where local activation of TRPM8 channels in sensory neurons mitigates the painful symptoms of this disease.

## Introduction

Benzodiazepines are a very well-studied class of drugs. They exert inhibitory effects on central nervous system excitability underpinning their clinical use as anxiolytics, hypnotics and anti-seizure medications. Their mechanism of action through allosteric modulation of central GABA_A_ receptors to enhance GABAergic tone^1,2^ has been well established over 50 years of study^3,4^. However, small molecule selectivity for a single class of target is rarely absolute, with the broader polypharmacological profile of any drug being critical for understanding physiological activity as well as unwanted side effects.

Here, we examined the polypharmacology of benzodiazepines in the context of engagement of transient receptor potential (TRP) ion channels. This interest was prompted by our recent discovery that clonazepam, a ‘first generation’ benzodiazepine^5^, activates a schistosome transient receptor potential melastatin (TRPM) ion channel^6,7^. Schistosomes are parasitic flatworms and the causative agents of the neglected tropical disease schistosomiasis (‘Bilharzia’). Despite the evolutionary distance between humans and platyhelminths, we were curious whether a similar profile of pharmacological regulation could exist in humans, underpinning the rationale for this study.

## Results

Clonazepam and meclonazepam (methyl-clonazepam) were recently shown to activate a schistosome transient receptor potential melastatin ion channel named TRPM_MCLZ_^6,7^. TRPM_MCLZ_ harbors a transmembrane ligand binding pocket that closely resembles the menthol binding pocket in TRPM8^6,7^. On the basis of this binding pocket homology, we screened a benzodiazepine library against human TRPM8 (hTRPM8) using an intracellular Ca^2+^ reporter assay. Whereas none of the screened benzodiazepines triggered changes in cytoplasmic Ca^2+^ in untransfected HEK293 cells, multiple analogs evoked Ca^2+^ signals in hTRPM8 expressing HEK293 cells (**Figure 1A**). The top three ‘hits’ in terms of peak amplitude were all 7-nitrobenzodiazepines: clonazepam, flunitrazepam and N-desmethylflunitrazepam (a metabolite of flunitrazepam, Supplementary Table 1).

**Figure 1.**
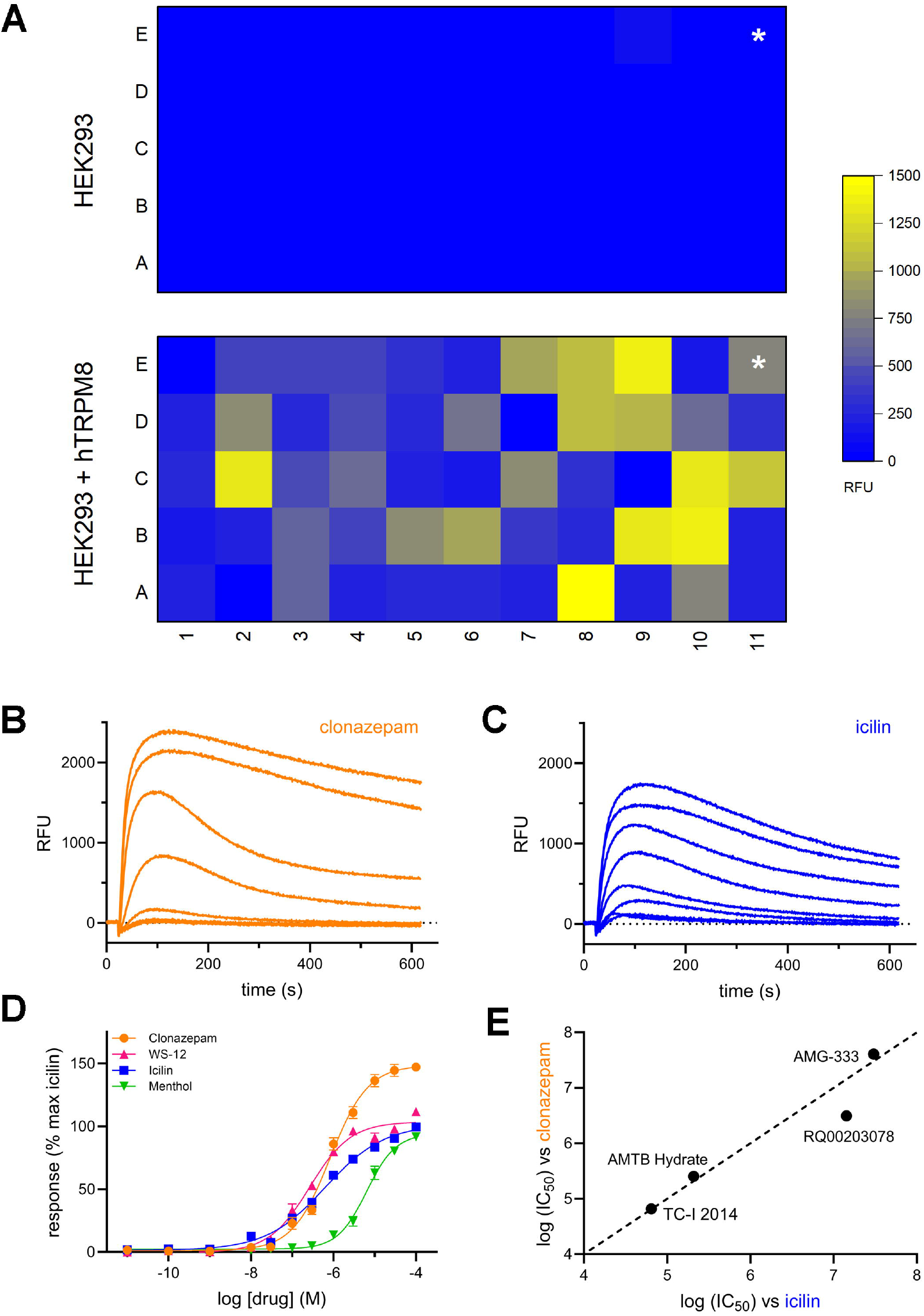
Benzodiazepine activation of human TRPM8. (**A**) Heatmap of Ca^2+^ signal amplitude (RFU, relative fluorescence units) evoked by 53 different benzodiazepines (A1 through to E9) together with icilin (E11, asterisked). Each benzodiazepine ligand was tested at ∼20 μM concentration and values are averaged from two biological replicates. A list of all the tested ligands is provided in Supplementary Table 1. (**B&C**) Example traces of fluo-4 NW fluorescence representing Ca^2+^ signals evoked by various concentrations (10nM to 100µM) of (B) clonazepam and (C) icilin in hTRPM8-expressing HEK293 cells. Data are plotted over same ranges. Also see Figure S1 & S2. (**D**) Concentration response curves for clonazepam, resourced from a different vendor, together with the canonical hTRPM8 agonists (icilin, WS-12 and menthol). Data represent mean ± SEM from n=3 independent experiments. Data are normalized to the peak response seen with icilin. (**E**) Plot of IC_50_ values for four different hTRPM8 antagonists in response to stimulation by clonazepam (y-axis) or icilin (x-axis). Also see Figure S1.

This same assay was then repeated with different lots of repurchased clonazepam to confirm activity at hTRPM8 (**Figure 1B**). Clonazepam elicited a concentration dependent activation of hTRPM8 (EC_50_=828±84nM), with the evoked Ca^2+^ signals displaying a greater peak magnitude than observed with icilin (**Figure 1C**), a Ca^2+^ dependent hTRPM8 agonist^8^, and other canonical hTRPM8 activators (**Figure 1D**). Robust Ca^2+^ signals with clonazepam were also apparent using Ca^2+^ reporter dyes of lower affinity (see Figure S1A & S1B). Finally, different hTRPM8 blockers were examined for effectiveness at attenuating clonazepam-evoked Ca^2+^ signals. Four hTRPM8 antagonists (TCI-2014, RQ00203078, AMTB and AMG-333) blocked clonazepam and icilin-evoked Ca^2+^ signals, with a similar rank order of potency (**Figure 1E**). In contrast, flumazenil, a benzodiazepine blocker of GABA_A_ responses^9^, had no effect on clonazepam-evoked Ca^2+^ signals (≤100µM, Figure S1C).

Electrophysiology was used as an orthologous method to investigate clonazepam action at hTRPM8. ‘Whole cell’ currents in HEK293 cells overexpressing hTRPM8 showed outward rectification in response to clonazepam and other hTRPM8 activators (see Figure S2A). The clonazepam-evoked inward current density was greater with clonazepam than with icilin or menthol (see Figure S2B). In response to cooling, clonazepam caused a shift of hTRPM8 responsiveness toward warmer temperatures, with half-maximal open probability occurring at 26.3±2.1□C in the presence of clonazepam, compared with ∼15.9±1.8□C in the absence of ligand (see Figure S2C & S2D). This behavior was similar to that seen with other hTRPM8 agonists, including WS-12 (Figure S2C & S2D) and menthol^10^.

Activity at other vertebrate TRPM8 channels was then examined. Clonazepam activated mouse TRPM8 (*Mm*.TRPM8, EC_50_=367±55nM) and rat TRPM8 (*Rn*.TRPM8, EC_50_=357±55nM), underscoring retention of benzodiazepine sensitivity at TRPM8 in commonly used animal models (**Figure 2A**). However, clonazepam did not activate an avian TRPM8 ortholog (*Ficedula albicollis* TRPM8, *Fa*.TRPM8, Figure 2A). These results parallel data obtained with the hTRPM8 agonist icilin, which lacks activity at avian TRPM8^8^. Responsiveness of avian TRPM8 to icilin is however restored when a human amino acid variant found within the third transmembrane helix (TM3) is engineered into the avian sequence^8^. Reciprocally, icilin sensitivity of hTRPM8 is lost when the avian variant is introduced into the hTRPM8 sequence^8^. Reciprocal mutants of the human and flycatcher channel were therefore profiled against clonazepam. Clonazepam activated the mutant avian construct, *Fa*.TPRM8[A796G] which harbors the mammalian amino acid variant at this position (**Figure 2B**), but not the hTRPM8 mutant, hTRPM8[G805A], where the avian amino acid variant was introduced into the hTRPM8 channel (Figure 2B). These data suggest commonalities in the binding mode of both ligands to the TRPM8 orthologs. Indeed, cheminformatic profiling of hTRPM8 ligands clustered clonazepam together with icilin relative to other commonly used hTRPM8 activators (**Figure 2C**). Finally, TRP channel selectivity was examined by screening clonazepam against other human TRP channels. No activation of other human TRP channels by clonazepam was observed (**Figure 2D**). Collectively, these data demonstrate clonazepam selectively and robustly activates mammalian TRPM8.

**Figure 2.**
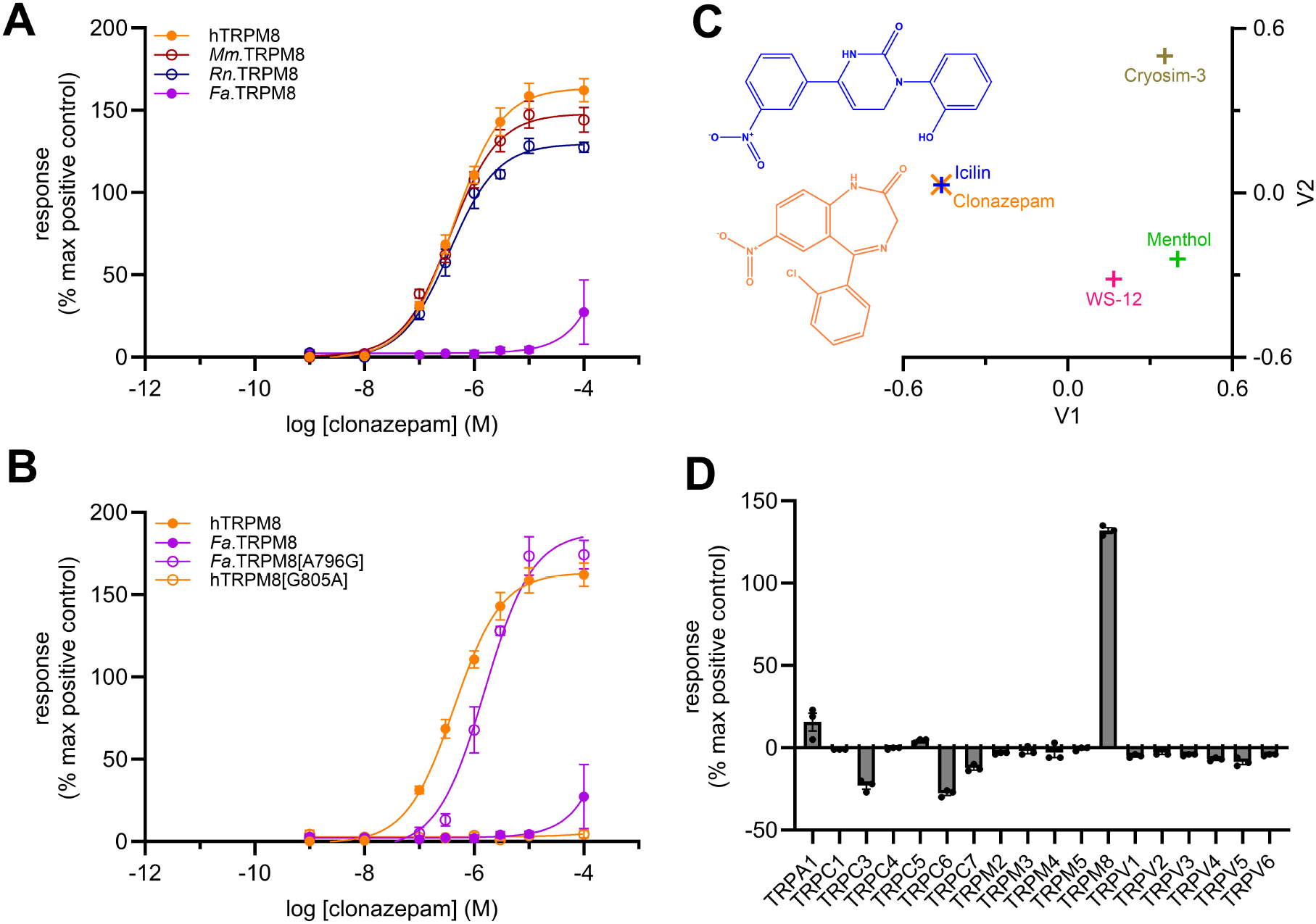
Activity of clonazepam at different TRPM8 ortholog and paralogs. (**A**) Concentration response curves in a Ca^2+^ reporter assay showing activation of mouse TRPM8 (*Mm*.TRPM8), rat TRPM8 (*Rn*.TRPM8), collared flycatcher TRPM8 (*Fa*.TRPM8) by clonazepam. (**B**) Reciprocal mutagenesis of human (hTRPM8) and avian (*Fa*.TRPM8) constructs at a key TM3 residue controls clonazepam sensitivity. Data are shown for hTRPM8[G805A] (loss of sensitivity) and *Fa*.TRPM8[A796G] (gain of sensitivity). (**C**) Cheminformatic analysis of five different hTRPM8 agonist chemotypes visualized using a multidimensional scaling plot underscores the similarity between clonazepam and icilin. (**D**) Responses to clonazepam (100 µM) were resolved across a panel of eighteen human TRP channels. Individual assay details for the different channels are detailed in the methods. Data are normalized to the positive control ligand for each channel, and are represented as mean ± SEM.

To study the effects of clonazepam on native hTRPM8, assays were performed in G-402 cells, a kidney epithelial cell line that endogenously expresses hTRPM8. In Ca^2+^ reporter assays in G-402 cells, clonazepam triggered Ca^2+^ signals (EC_50_=597±64nM) which again were larger in magnitude than responses elicited by icilin or WS-12 (**Figure 3A**). In ‘cell attached’ recordings, clonazepam activated an outward rectifying, non-selective cation current in G-402 cells, similar to responses evoked by canonical hTRPM8 agonists (**Figure 3B**). These currents were blocked by hTRPM8 antagonists (see Figure S3A). Single channel responses evoked by clonazepam displayed a unitary conductance of 70±1pS (see Figure S3B & S3C), consistent with prior studies of hTRPM8^11^.

**Figure 3.**
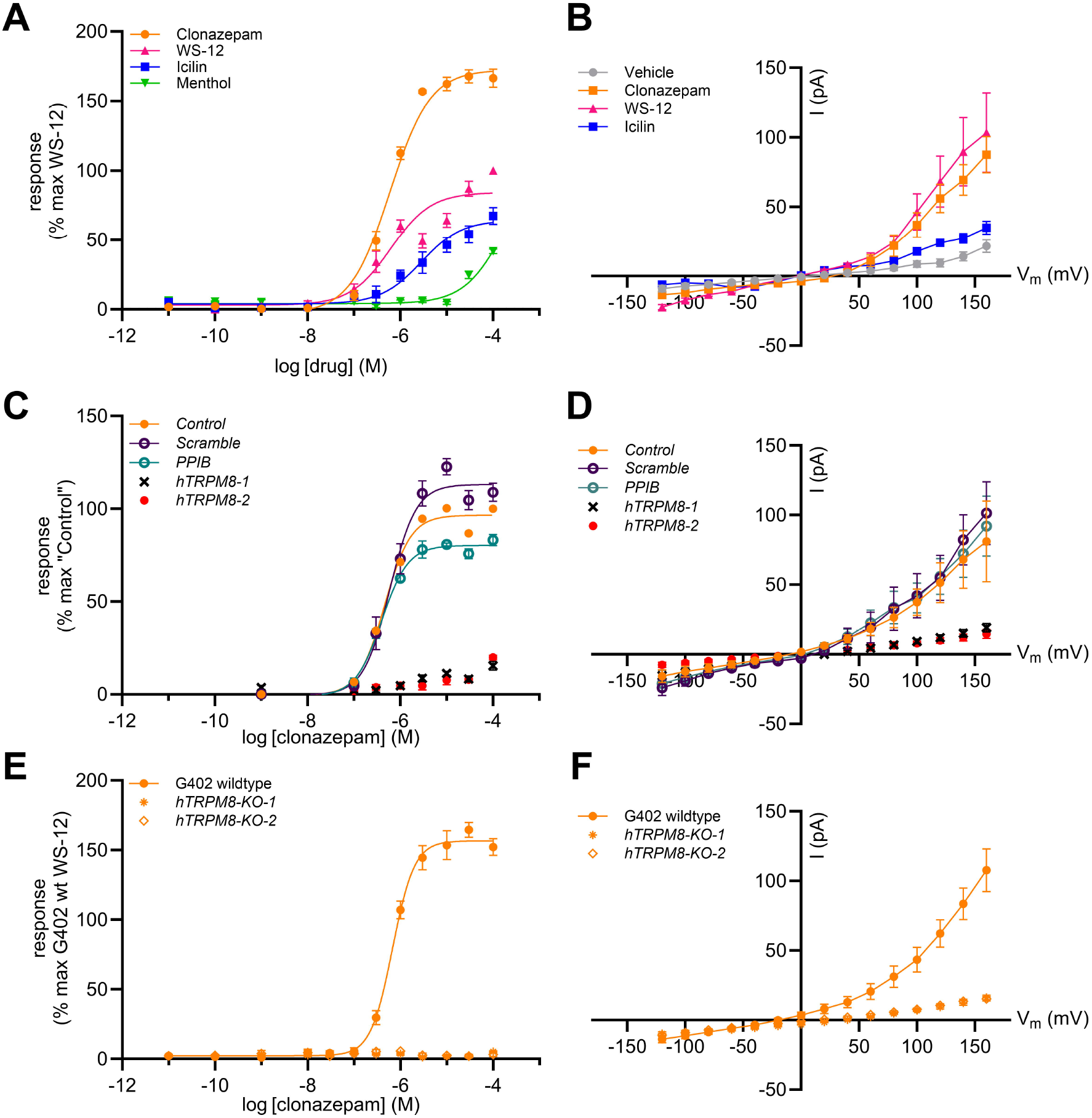
Clonazepam activation of native hTRPM8 channels. (**A**) Concentration response curves for clonazepam, icilin, WS-12 and menthol in wildtype G-402 cells resolved using a Ca^2+^ reporter assay. Data are normalized to the peak response seen with WS-12. (**B**) Current-voltage (I-V) plot in G-402 cells recorded in the ‘cell-attached’ mode in the absence (DMSO vehicle, 0.1%) or presence of various hTRPM8 agonists (each at 10 µM) in the pipette solution. Also see Figure S3. (**C**) Concentration response curves for clonazepam in G-402 knockdown cells resolved using a Ca^2+^ reporter assay and normalized to a ‘no RNA’ control (*Control*). The same assay is shown are shown for cells treated with dual control (*scramble, PPIB*) or hTRPM8-targeting siRNA constructs (*hTRPM8-1* or *hTRPM8-2*). Also see Figure S4. (**D**) Current-voltage (I-V) plot of G-402 cells in the ‘cell-attached’ mode that had previously been treated with the same siRNA constructs. Clonazepam (10 µM) was present in the pipette solution. (**E**) Concentration response curves for clonazepam evoked responses in G-402 wildtype and two clonal hTRPM8 knockout cell lines (*hTRPM8-KO-1* or *hTRPM8-KO-2*). Responses were normalized to the peak WS-12 response of G-402 wildtype cells. (**F**) Current-voltage (I-V) plot of G-402 knockout cell lines (*hTRPM8-KO-1* and *hTRPM8-KO-2*) recorded in ‘cell-attached’ mode in the presence of clonazepam (10 µM) in the pipette solution.

Endogenous hTRPM8 channels in G-402 cells were then targeted for knockdown using two different siRNA constructs (*hTRPM8-1, hTRPM8-2*). Knockdown of hTRPM8 using either construct inhibited Ca^2+^ signals evoked by clonazepam (**Figure 3C**). Clonazepam-evoked Ca^2+^ signals however persisted in G-402 cells treated with either of two control siRNA constructs (*scramble* or *PPIB*; Figure 3C). Similar results were seen in electrophysiological assays (**Figure 3D**). The effectiveness of each siRNA construct used for knockdown was confirmed using real-time PCR (see Figure S4). Finally, hTRPM8 knockout G-402 lines (*hTRPM8-KO-1, hTRPM8-KO-2*) were generated by CRISPR-Cas9 editing. In both these hTRPM8-KO cell lines, responses to clonazepam were not apparent, assessed either by Ca^2+^ reporter assay (**Figure 3E**) or electrophysiology (**Figure 3F**).

Clinically, clonazepam functions as an anti-seizure and anxiolytic drug via GABA_A_ receptor activation^2^. Plasma concentrations considered therapeutic for these effects (20-70ng/ml^12^, ≤200nM) are lower than those shown here to activate hTRPM8 (EC_50_ for clonazepam at hTRPM8 ∼600nM, Figure 2). However, the interaction of clonazepam with hTRPM8 could nevertheless have relevance for specific conditions. Most obvious is the disease known as burning mouth disorder (BMD^13^), an extremely painful chronic oral disorder hypothesized to result from sensory neuropathy^14,15^. Topical clonazepam yields relief from symptoms in most patients^16-18^, although there is no correlation between the efficacy of various benzodiazepines in treating BMD, systemic plasma concentration, and alleviation of anxiety or depressive symptoms^17,18^. The high, local concentration of clonazepam effective clinically for treating BMD (60-300µM in mouthwash, or similarly high local concentrations attained by sucking a sublingual tablet) surpasses the concentration range for clonazepam-evoked hTRPM8 activation. This suggests that the therapeutic action of clonazepam in BMD is likely realized by localized activation of hTRPM8 present within the sensory nerve pathways that innervate the mouth and tounge^19-21^. Assignment of hTRPM8 as the oral target of clonazepam would catalyze development of improved therapeutics for this condition.

To explore this idea, responses to clonazepam were examined in a primary culture of mouse trigeminal neurons. Trigeminal sensory neurons, which innervate the mouth and tongue, are functionally heterogeneous with multiple classes of cold sensing neurons defined physiologically or molecularly^22-24^. In Ca^2+^ imaging assays, a subset of isolated neurons exhibited responses to the TRPM8 agonist WS-12 (∼24% of total neurons, **Figure 4A**). Almost all of the WS-12 responsive neurons co-responded to clonazepam (∼95%, Figure 4A), with a Ca^2+^ signal that kinetically resembled the WS-12 response. To mitigate for any desensitization in this dual agonist application assay, responses were compared between trigeminal neurons exposed to a single application of either clonazepam or WS-12 in either wild type (**Figure 4B**), or TRPM8 knock out (*TRPM8*-KO) mice (**Figure 4C**). Ca^2+^ signals evoked by clonazepam were observed in wild type mice but were absent from *TRPM8*-KO mice (Figure 4B & 4C). Analysis of these Ca^2+^ signals revealed responses to clonazepam trended higher in magnitude and prevalence than with WS-12 (**Figure 4D & 4E**). Overall, these data demonstrate that clonazepam evokes TRPM8-dependent Ca^2+^ signals in trigeminal mouse neurons.

**Figure 4.**
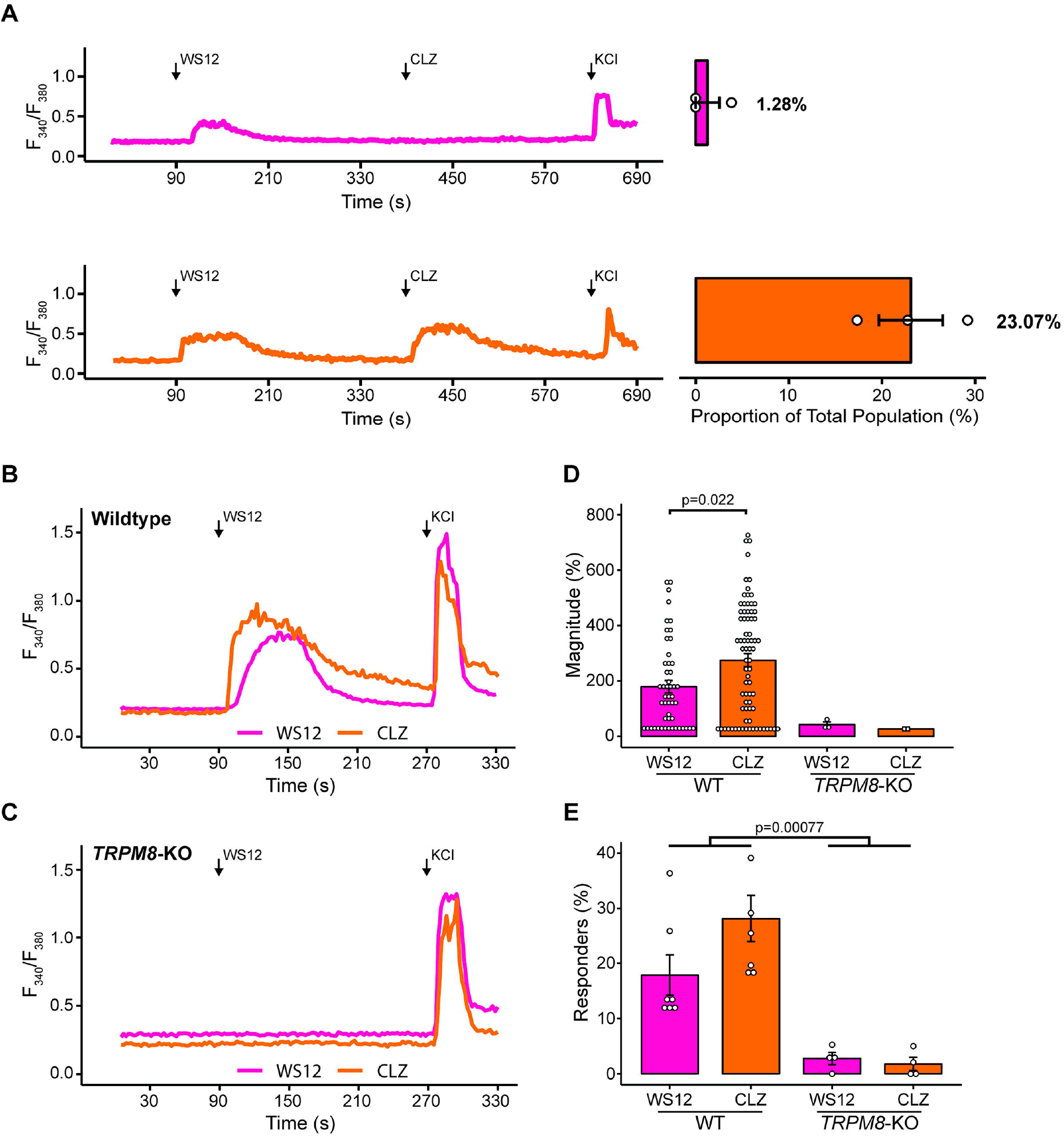
Clonazepam evokes Ca^2+^ signals in TRPM8-positive trigeminal neurons. (**A**) Representative traces of fura-2 fluorescence ratio (F_340_/F_380_) over time in WS-12 responding neurons that were exposed first to WS-12 (10 μM) and then to clonazepam (10 μM). Neurons were classified as single responders (to WS-12 alone, pink) or as dual responders (WS-12 and clonazepam, orange). The proportion (percentage of total neurons in each imaging field) of each class of neuron is shown on the right. (**B & C**) Comparison of responses to application of WS-12 (10 μM) or clonazepam (10 μM) in (B) wildtype or (C) *TRPM8*-KO neurons. These data are quantified in terms of (**D**) signal amplitude and (**E**) number of cells responding. A total of ten (n=2 males, n=8 females) C57BL/6 mice and four (n=2 males, n=2 females) *TRPM8* knockout mice were used for these experiments, with 22-55 neurons imaged per mouse.

In conclusion, this report demonstrates that clonazepam, as well as several other 7-nitrobenzodiazepines, activate hTRPM8. This discovery provides new insight into benzodiazepine polypharmacology and highlights opportunity to explore the benzodiazepine scaffold for hTRPM8 modulation. Further, this result identifies a local mechanism of clonazepam action to mitigate the painful symptoms of BMD.

## Author Contributions

Conceptualization: LE, JSM; Formal analysis LE, EGC, JE; Investigation: LE, EGC, JE, ADM, SKP; Project administration: CLS, JSM; Supervision: JSM; Funding acquisition: JSM; Writing original draft: LE, JSM; Writing, review & editing: all authors.

## Acknowledgements

Work was supported by NIH (NS070711 to CLS; AI145871 to JSM). LE was supported by a Graduate Fellowship from the Medical College of Wisconsin Cancer Center. JE was supported by a Post-doctoral Fellowship (F32NS138223-01).

## EXPERIMENTAL MODEL AND STUDY PARTICIPANT DETAILS

### Mouse breeding and housing

C57BL/6 mice (JAX: 000664) and TRPM8 knockout mice (B6.129P2-Trpm8tm1Jul/J, JAX: 008198) were procured from Jackson Labs. All mice were housed within Medical College of Wisconsin (MCW) facilities according to NIH stipulated guidelines. All animal protocols were approved by the Institutional Animal Care and Use Committee at MCW.

### Cell lines and reagents

HEK293 cells (ATCC CRL-1573, female) were cultured in high glucose, L-glutamine, Phenol Red, Sodium pyruvate Dulbecco’s modified Eagle’s medium supplemented (DMEM, Thermo Fisher Scientific, Cat#11995073) with 10% fetal bovine serum (FBS, Thermo Fisher Scientific, Cat#A5256701), penicillin (100 units/ml), streptomycin (100 μg/ml), and L-glutamine (290 μg/ml) (Thermo Fisher Scientific, Cat#10378016). G-402 cells (ATCC CRL-1440, female) were cultured in McCoy’s 5A Medium (Thermo Fisher Scientific, Cat#16600082) supplemented with 10% FBS, penicillin (100 units/ml), streptomycin (100 μg/ml), and amphotericin B (0.25 μg/ml) (Thermo Fisher Scientific, Cat#15240062). All cell lines were cultured in a humid environment at 37°C and 5% CO_2_. All lines were tested monthly for Mycoplasma contamination via PCR (Millipore Sigma, Cat#MP0035) and quarterly via antigen (Lonza Bioscience, Cat# LT07-318). All cell lines, including engineered lines, were authenticated by short tandem repeat DNA profiling (ATCC).

### *Ex* vivo trigeminal nerve harvest and culture

To isolate trigeminal ganglion (TG) neurons, mice were deeply anesthetized by isoflurane inhalation prior to euthanasia by decapitation. Whole TG were then incubated in collagenase IV (1 mg/mL) in DMEM/F12 (Thermo Fisher Scientific, Cat#21331020) for 45 minutes at 37°C. Ganglia were then incubated in 0.05% trypsin (Thermo Fisher Scientific, Cat# 25300062) in DMEM for 45 minutes at 37°C, which was immediately thereafter neutralized with horse serum (Thermo Fisher Scientific, Cat#26050088). Ganglia were then washed and resuspended in complete media (DMEM/F12, 10% inactivated fetal horse serum, 1% glucose, 2 mM L-glutamine, 100 U/mL penicillin, and 100 µg/mL streptomycin). Ganglia were progressively mechanically dissociated with a p1000 and p200 micropipette prior to plating on laminin-coated 12 mm round glass coverslips. Dissociated neurons were fed with complete media 1-2 hours after plating. Dissociated TG neuronal somata were used for calcium imaging 18-24 hours after plating.

## METHOD DETAILS

### Chemical preparation

hTRPM8 control ligands (icilin, WS-12, and menthol) were sourced from Cayman Chemical and solubilized in 100% DMSO (Sigma-Aldrich). The benzodiazepine library was procured from Cayman Chemical, solubilized in 135µL of DMSO (final concentration ∼20 µM).

### Plasmid preparation

cDNA constructs for TRPM8 orthologs were cloned in a pcDNA3.1 backbone sourced as: *Homo sapiens* TRPM8 *(hTRPM8;* NCBI: NM_024080.5, UniProt: Q7Z2W7), *Mus musculus* TRPM8 (*Mm*.TRPM8; NCBI: NP_599013.1, UniProt: Q8R4D5), and *Ficedula albicollis* (*Fa*.TRPM8; NCBI: XP_005049003.1) by GenScript. *Rattus norvegicus* TRPM8 (*Rn*.TRPM8) tagged with GFP^25^ was a gift from Sebastián Brauchi (Addgene plasmid #64879; RRID:Addgene_64879).

### Over-expression studies

Transient transfections of HEK293 cells were performed using Lipofectamine 2000, following the manufacturer’s protocol (Thermo Fisher Scientific). For a 60mm petri dish, cells (seeded at 1×10^6^ cells) were incubated with OptiMEM (800µl) containing a mixture of lipofectamine 2000 (20µl) and DNA (8μg).

### Knockdown studies

Endogenous expression of hTRPM8 in G-402 cells was ablated using gene-specific siRNA constructs transfected using Lipofectamine 3000 (Thermo Fisher) following the manufacturer’s protocol. The initial seeding densities of wildtype cells were 1.5×10^5^, 5×10^5^, 8.5×10^5^ cells in 35mm, 60mm, and 100mm dishes, respectively. Dual siRNA constructs targeting either exon 5 (Silencer® siRNAs, Cat No. AM16708, ID 104796, “*hTRPM8-1”)* or exon 26 (Silencer® siRNAs, Cat No. AM16708, ID 30886, “*hTRPM8-2”*) were used to knockdown hTRPM8. An anti-cyclophilin B (*PPIB*) siRNA (Cat. No. AM16078, ID: 110793), Silencer™ Negative Control No. 1 siRNA (*Scramble*) (Cat. No. AM4611, Invitrogen), and water were used as controls. Dishes were incubated in siRNA-containing medium (33pmol siRNA/mL) for no more than 4.5 hours, after which the medium was aspirated and replaced with growth medium containing no antibiotics. Cells were re-plated on glass chips for electrophysiology or directly into FLIPR plates for the high-throughput Ca^2+^ reporter assay.

### Generation of TRPM8 knockout cell lines

The G-402 hTRPM8-knockout cell lines were generated by Applied Biological Materials Inc. (Vancouver, Canada). A lentiviral and CRISPR-Cas-9 package with three unique hTRPM8 sgRNA targets (hTRPM8 sgRNA#1-3) were transfected into wildtype cells. Cells containing the lentiviral construct were selected using puromycin treatment and serially diluted to select single clones for expansion. Validation of knockout was confirmed by Sanger sequencing using the following forward and reverse primers (TRPM8#1F and TRPM8#1R). Two clones *hTRPM8-KO-1* and *hTRPM8-KO-2*) were confirmed to harbor biallelic heterozygous frameshift mutations.

### Quantitative PCR

RNA was isolated using the RNeasy Mini Kit according to the manufacturer’s protocol (Qiagen). Equivalent amounts of cDNA (1µg) were generated using a High-Capacity RNA-to-cDNA™ Kit (Applied Biosystems) and diluted 1:5 v/v for PCR analyses. Primers for qPCR were synthesized by Integrated DNA Technologies. Quantitative PCR reactions were performed as 10µL reactions in 96-well plates using 8µL of PowerUp™ SYBR™ Green Master Mix supplemented with 3µM of gene specific primers (3µM) and 2 µL of diluted cDNA. Plates were sealed and briefly spun down prior to PCR (CFX Duet Real-Time PCR System, BioRad). The PCR protocol was as follows: uracil-DNA glycosylase degradation 50°C for 2 mins, hot-start activation at 95°C for 2 mins, denature 95°C for 15 s, anneal 59°C for 15s, and 72°C for 1 min, repeated for 40 cycles. Bio-Rad CFX manager was used to track individual well RFUs. A quantification cycle (Cq) threshold of 200 RFUs was set for all experiments. Values were normalized to GAPDH expression using the ΔΔCt method. Data were visualized using GraphPad Prism (v.10.6.1).

### Calcium reporter assay

A Ca^2+^ reporter assay using a Fluorescence Live Imaging Plate Reader assay system (Molecular Devices, FLIPR^TETRA^) was used for functionally profiling TRPM8. HEK293 cells were seeded (20,000 cells/well) in a black wall, clear bottom, poly-L-lysine coated 384-well plate and incubated (16-24 hours) in DMEM supplemented with 10% FBS. G-402 cells lines were seeded (13,500 cells/well) and incubated (40-48 hours) in McCoy’s 5A medium + 10% FBS. Prior to the assay, cells were incubated in 20 µL assay buffer (HBSS + 20 mM HEPES at pH 7.4, supplemented with probenecid (2.5 mM) and the fluorescent Ca^2+^ indicator Fluo-4NW (Thermo Fisher Scientific, Cat#F36206)) for 45-50 min at 37°C and 15 min at room temperature. Drug plates were prepared using assay buffer without probenecid or fluorescent dye. After dye loading, the FLIPR assay was performed by measuring basal fluorescence of each well for 20s prior to drug addition (5 µL of 5x drug stock). Raw-fluorescence was captured for another 250-600s following drug addition. For antagonist assays, antagonists (5 µL of 6x drug stock) were added with fluorescence monitored for 250s prior to subsequent injection of 1µM icilin or CLZ (5µL of 6µM stock) with fluorescence continually monitored for a further 250-600s. For assays with lower Ca^2+^ affinity fluorescence dyes (Fluo-4FF, AM and Fluo-5F, AM, Thermo Fisher Scientific, Cat#F23980 and Cat#F14221), an additional wash step (HBSS with 20 mM HEPES, pH = 7.4) was included after dye loading. All assays were performed at room-temperature and included vehicle, positive (icilin and/or WS-12), and negative (buffer, and/or non-transfected cells) controls. Peak fluorescence signals (relative fluorescence units, RFU) were corrected for baseline and normalized to relevant positive controls (100µM icilin or WS-12). For concentration response curves, zero was defined as the smallest value within a given biological replicate.

### Cheminformatic analysis

TRPM8 agonist compound information was entered into the ChemMine Tools web server^26^ and clustered in a 2D-MDS with a similarity cutoff of 0.4 following the approach of Mebrat *et al*^27^.

### Human TRP channel profiling

Clonazepam was profiled against a panel of 18 human TRP channels (SB Drug Discovery, Glasgow). In these assays, HEK293 cells expressing specific human TRP channels were challenged with clonazepam and responses resolved using either a membrane potential dye (TRPC1, TRPC3, TRPC4, TRPC5, TRPC6, TRPC7, TRPM4 and TRPM5; Molecular Devices, R8126) or a Ca^2+^ reporter (TRPA1, TRPV1, TRPV2, TRPV3, TRPV4, TRPV5, TRPV6, TRPM2, TRPM3 and TRPM8; Calcium 5, Molecular Devices, R8186) based on the ionic permeability of each channel. Responses were evaluated relative to peak responses obtained with reference compounds (ASP7663, TRPA1; Englerin A, TRPC1, TRPC4; GSK-1702934A, TRPC3, TRPC6; Carbachol, TRPC5, TRPC7; H_2_O_2_, TRPM2; CIM0216, TRPM3; ionomycin, TRPM4, TRPM5; icilin, TRPM8; capsaicin, TRPV1; cannabidiol, TRPV2; 2-APB, TRPV3; RN1747, TRPV4; CaCl_2_, TRPV5, TRPV6). For each assay, cells were trypsinised, counted and seeded in black, clear bottomed 384 well plates (10,000 cells per well) and incubated overnight. Media was then removed from cell plates and assay buffer (30 μl) supplemented with reporter dye (10 μl) added prior to incubation (37□C, 40 mins). Plates were then placed in a FLIPR (Molecular Devices, serial no: FX01138), and fluorescence monitored (488nm/510-570nm) every second prior to addition of clonazepam (10µM) after which recordings continued for 5 minutes at room temperature

### Electrophysiological recording

Cells were plated onto glass coverslips 24-48 h prior to being secured in an Olympus BX51WI inverted microscope recording chamber. A multiClamp 700B amplifier and Digidata 1440A digitizer (Molecular Devices, San Jose, CA) were used for electrophysiological recordings. Signals were passed through an 8-pole Bessel low pass filter at 1 kHz and sampled at 10 kHz. Data analysis was performed using Clampfit 11 software (Molecular Devices). Pipettes were pulled from borosilicate glass (BF150-110-10, Sutter Instruments, Novato, CA) to a resistance of 6-8 MΩ. For ‘cell-attached’ recordings, both pipette and bath solutions were Hank’s buffered saline solution (HBSS) supplemented with HEPES (20mM). In ‘whole cell’ and ‘outside-out’ recordings the pipette solution comprised CsCH_3_SO_3_ (140mM), CsCl (5mM), HEPES (10mM), EGTA (1mM), at pH 7.4 (with CsOH). Holding membrane voltage (V_m_) was -80 mV. All recordings were performed at room temperature except recordings in ‘outside-out’ mode that were performed across a temperature range ∼8-42□C. To estimate hTRPM8 temperature dependence 160 ms long voltage ramps from -80 to 80 mV were applied to assess channel activity at different temperatures. Ligands were applied to the bath solution at 10 µM and DMSO (0.1%) used as a control. Temperature was controlled by a temperature controller TC-324B (Warner Inst.). After formation of an ‘outside-out’ patch, the recording chamber was perfused with cooled solution (∼6-8□C) and then heated up to ∼40-42□C. Open probability (P_open_) was estimated as normalized peak current at 80 mV (I/I_max_) plotted against temperature. Typically, P_open_ was steady at ∼1 at cooler temperatures and approached 0 at warmer temperatures. The temperature at which the P_open_= 0.5 was defined as ET_50_. Tukey’s test was used to compare ET_50_ values between control and ligand treated conditions.

### Calcium imaging of primary mouse neurons

Somata were washed with extracellular normal HEPES (ENH) buffer (in mM: 150 NaCl, 10 HEPES, 8 glucose, 5.6 KCl, 2 CaCl_2_, 1 MgCl_2_; pH 7.4 +/-0.03, mOsm 320 +/-3) for 1-2 minutes, incubated with 2.5 mg/mL Fura-2AM (Life Technologies, Cat#F1221) in 2% bovine serum albumin for 45 minutes at room temperature, and washed again in ENH at room temperature for 30 minutes. Coverslips were then mounted to an imaging stage and perfused with ENH (6 mL/min). Fura-2 fluorescence was measured with excitation at 340 nm and 380 nm and captured by a cooled DS-Qi1MC monochrome camera (Nikon). NIS Elements software (Nikon) was used to analyze signals following stimulation with 340 nm or 380 nm and calculated bound/unbound calcium ratio (340/380). To induce intracellular calcium transients, neuronal somata were exposed to WS12 (10 µM, 1 minute), or clonazepam (10 µM, 1 minute) followed by a 2-6 minute washout in ENH depending on the experiment. Cell viability was assessed by transient (20 second) exposure to 50 mM KCl followed by washout with ENH. Somata exhibiting at least 20% increase in 340/380 ratio relative to most recent baseline or washout were considered positive responders. Cells were only included for analysis if they exhibited no positive response to initial ENH exposure and exhibited a positive response to at least one test compound or 50 mM KCl.

## QUANTIFICATION AND STATISTICAL ANALYSIS

All HEK293 experiments comprised at least three biological replicates (independent transfections). Within each of these assays, individual analogs or constructs were screened in triplicate (technical replicates). All G-402 experiments shown contained at least three replicates, with biological replicates defined as unique plates. Data in concentration response curves were normalized with zero defined as the smallest value within a replicate and 100 was defined as the peak icilin or WS-12 response. Curve fitting was performed using non-linear dose-response algorithm in GraphPad Prism (v.10.6.1). All electrophysiology experiments shown were from a minimum of three unique recordings. Data were analyzed using Clampfit 11 (v.4) and visualized using Origin (v.10.2.0.188). Sex of mice included in given TG experiments are located in figure legends. No significant difference between male and female mice were observed. Statistical analyses for TG calcium imaging experiments were performed using the R programming language (version 4.3.2) and the RStudio environment. Comparisons of response and co-response frequency considered results from individual mice as samples, treated test-compound exposure as a within-sample independent variable, and thus used a paired t-test. Response magnitude was compared at an individual cell level, precluding the use of within-sample statistics. As these results violated the assumption of normality, they were compared with the Mann-Whitney U test. For Figure 2E, a 2-way ANOVA with repeated measures was used, treating “Agonist” as the within-subjects independent variable and “Genotype” as the between-subjects independent variable.

## KEY RESOURCES TABLE

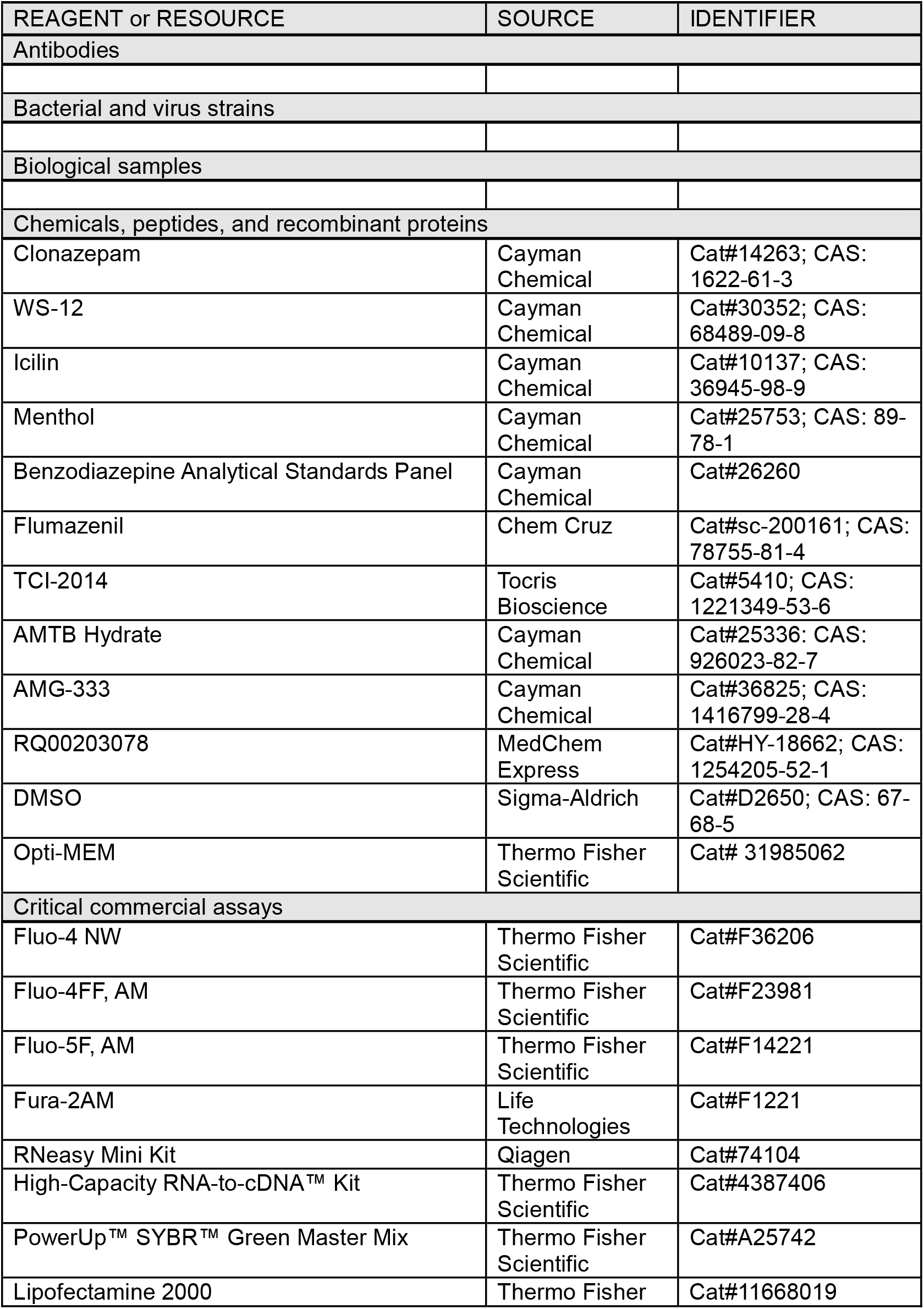
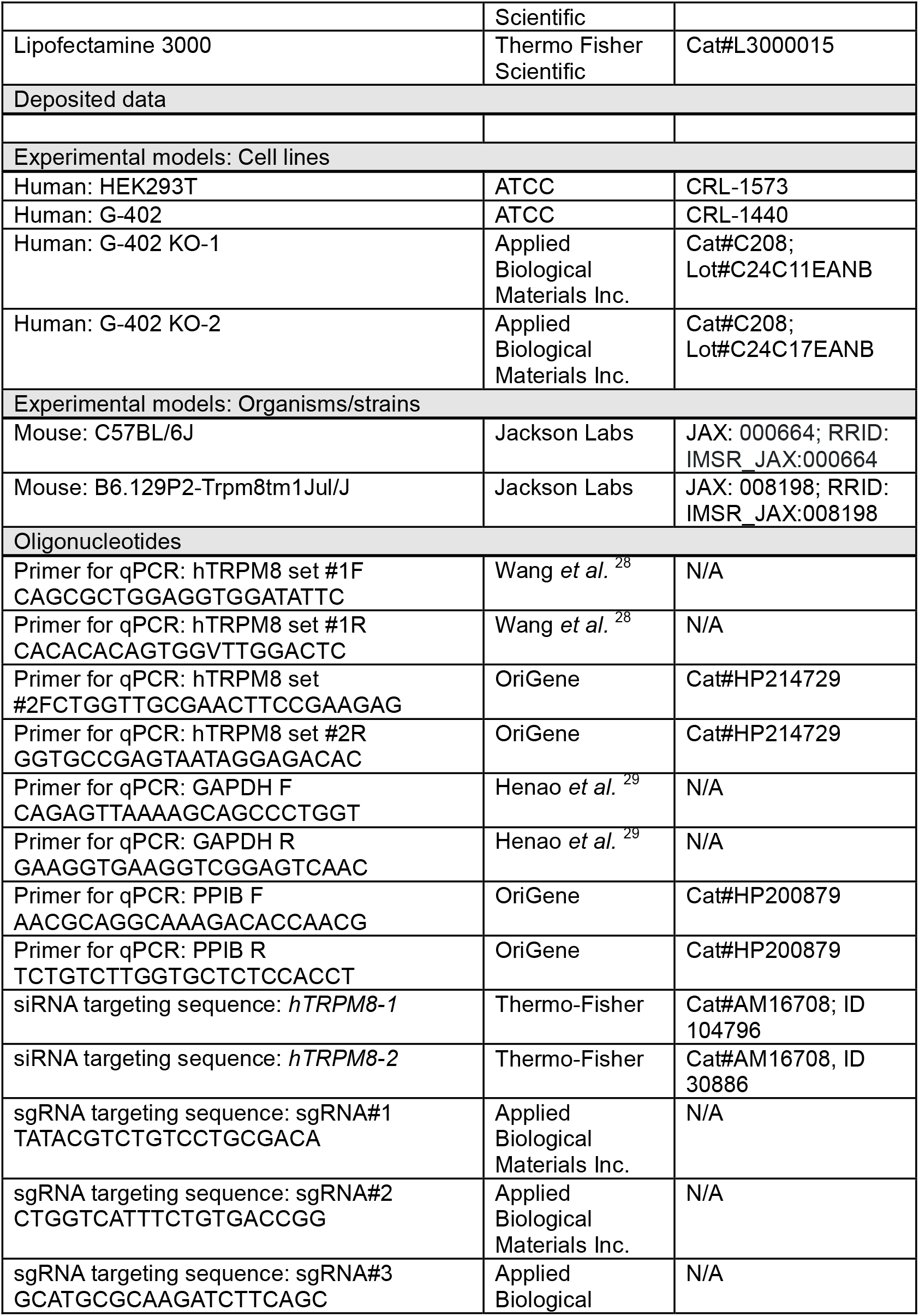
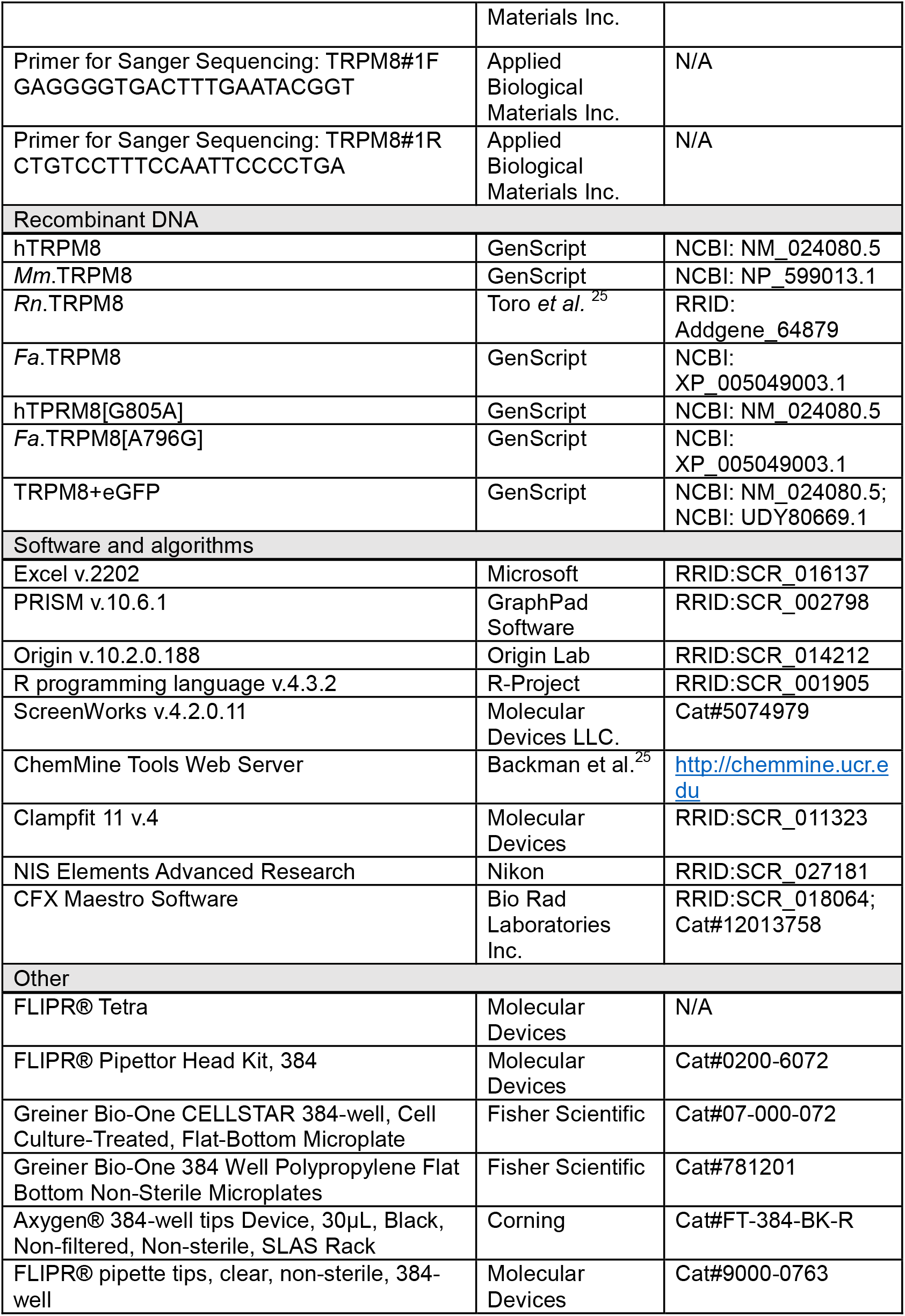
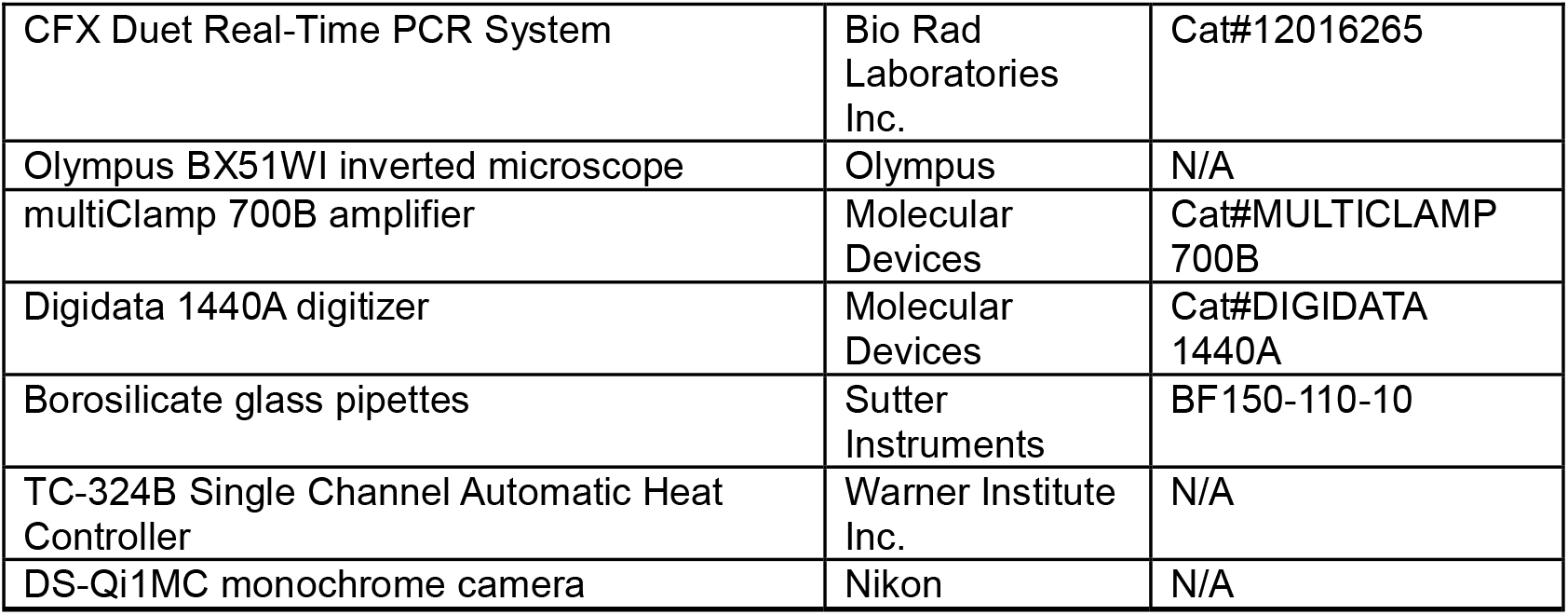

