## Supplementary Material for "Clonazepam activates the transient receptor potential melastatin 8 (TRPM8) ion channel"

**Supplementary Table 1. Results from benzodiazepine library screen, related to Figure 1A.** Table lists 53 benzodiazepines by location on plate stratified by peak amplitude of $Ca^{2+}$ signal shown normalized relative to icilin. Ligands are grouped as 1st or 2nd generation benzodiazepines.

**Supplementary Figure 1. Clonazepam-evoked $Ca^{2+}$ signals resolved with different fluorescent $Ca^{2+}$ indicators, related to Figure 1.** (**A & B**) Concentration response curves for clonazepam, icilin, and menthol in hTRPM8-expressing HEK293 cells reported using (A) Fluo-5F ($K_d$ for $Ca^{2+}$ ~2.3μM) and (B) Fluo-4FF ($K_d$ for $Ca^{2+}$ ~9.7μM). Data are normalized to the peak response seen with icilin and represent mean±SEM from n=3 independent experiments. (**C**) Effect of flumazenil on clonazepam, icilin, WS-12 and menthol evoked $Ca^{2+}$ signals. Each agonist was delivered at an ~$EC_{80}$ concentration and peak $Ca^{2+}$ signal amplitude resolved in the presence of increasing concentrations of flumazenil. Data are normalized to peak agonist response in the absence of flumazenil and show mean±SEM from n=3 independent experiments.

**Supplementary Figure 2. Electrophysiological analysis of clonazepam and TRPM8 agonists in hTRPM8 transfected HEK293 cells, related to Figure 1.** (**A**) Normalized current-voltage plot of the 'whole-cell' current in hTRPM8 transfected HEK293 cells in the presence of clonazepam (5 μM), icilin (5 μM), or menthol (500 μM) in the bath solution (HBSS). (**B**) Comparison of current density (I, pA/pF) of the 'whole-cell' current at -80 mV in hTRPM8 transfected HEK293 cells in the presence of clonazepam (5 μM), icilin (5 μM), or menthol (500 μM) in the bath solution. Significance between cohorts was assessed using Tukey's test (mean±SEM; ***, P<0.001). (**C**) Effect of cooling on hTRPM8 responsiveness to clonazepam. Scatterpolt shows the effect of temperature on hTRPM8 open probability in the absence (Ctrl, vehicle DMSO 0.1%) or presence of either CLZ or WS-12 (10 μM) in the bath solution. (**D**) Bar graph depicting measurements of temperature at which hTRPM8 displayed half-maximal open probability ($ET_{50}$). Data are shown as mean ± SEM with significance assessed using Tukey's post hoc t-test, *** p<0.01.

**Supplementary Figure 3. Electrophysiological analysis of hTRPM8 in G402 cells, related to Figure 2.** (**A**) Effect of various TRPM8 antagonists on clonazepam-evoked $Ca^{2+}$ currents in G402 cells. Representative current-voltage recordings made from G402 cells in 'cell-attached' mode in the presence of clonazepam (10 μM), or clonazepam with AMTB or TC-I 2014 (each at 50 μM) in the bath solution. Holding voltage was -80 mV. (**B**) Single-channel traces recorded from G402 cells in 'cell-attached' mode at 40 or 60 mV in response to clonazepam (10 μM) added to the pipette solution. 'c', closed state. (**C**) Slope conductance of single-channel currents obtained by linear regression, estimating single-channel conductance at 70±1pS (mean±SEM, n=3).

**Supplementary Figure 4. Quantifying penetrance of knockdown in G402 cells, related to Figure 3C & 3D.** Quantitative PCR of G402 cells for (**A**) hTRPM8 and (**B**) cyclophilin B (PPIB) expression. Cq values were normalized to GAPDH and G402 *'Control'* using the $\Delta\Delta C_t$ method. A two-way ANOVA with a Dunnett's test was performed illustrated with *, P<0.05; **, P<0.005; ****, P<0.0001.

| Item name | Molecular Formula | HEK293 + hTRPM8 | % icilin | Generation | library # | Row&Column |
|---|---|---|---|---|---|---|
| **Clonazepam** | **C15H10ClN3O3** | **1478.52** | 185.10 | **1** | **8** | **A8** |
| N-Desmethylflunitrazepam | C15H10FN3O3 | 1381.93 | 173.01 | 1 | 53 | E9 |
| Flunitrazepam | C16H12FN3O3 | 1355.05 | 169.64 | 1 | 21 | B10 |
| Desalkylflurazepam | C15H10ClFN2O | 1337.93 | 167.50 | 1 | 32 | C10 |
| Delorazepam | C15H10Cl2N2O | 1320.88 | 165.37 | 1 | 20 | B9 |
| Nitrazepam | C15H11N3O3 | 1303.14 | 163.15 | 1 | 24 | C2 |
| N-Desmethylclobazam | C15H11ClN2O2 | 1130.38 | 141.52 | 1 | 33 | C11 |
| Phenazepam | C15H10BrClN2O | 1065.95 | 133.45 | 1 | 52 | E8 |
| Difludiazepam | C16H11ClF2N2O | 1011.24 | 126.60 | 1 | 42 | D9 |
| Nordiazepam | C15H11ClN2O | 964.64 | 120.77 | 1 | 17 | B6 |
| Desmethylclotiazepam | C15H13ClN2OS | 950.36 | 118.98 | 1 | 51 | E7 |
| Nimetazepam | C16H13N3O3 | 818.07 | 102.42 | 1 | 29 | C7 |
| Fludiazepam | C16H12ClFN2O | 806.55 | 100.97 | 1 | 35 | D2 |
| Lorazepam | C15H10Cl2N2O2 | 802.62 | 100.48 | 1 | 16 | B5 |
| Flubromazepam | C15H10BrFN2O | 791.81 | 99.13 | 1 | 10 | A10 |
| Methylclonazepam | C16H12ClN3O3 | 652.07 | 81.64 | 1 | 39 | D6 |
| 3-hydroxy Phenazepam | C15H10BrClN2O2 | 585.69 | 73.32 | 1 | 3 | A3 |
| Diazepam | C16H13ClN2O | 574.07 | 71.87 | 1 | 14 | B3 |
| Ethyl Loflazepate | C18H14ClFN2O3 | 424.10 | 53.09 | 1 | 46 | E2 |
| Meclonazepam | C16H12ClN3O3 | 405.05 | 50.71 | 1 | 47 | E3 |
| Oxazepam | C15H11ClN2O2 | 376.45 | 47.13 | 1 | 18 | B7 |
| 7-Aminonimetazepam | C16H15N3O | 319.02 | 39.94 | 1 | 30 | C8 |
| Temazepam | C16H13ClN2O2 | 297.79 | 37.28 | 1 | 19 | B8 |
| 7-Aminoflunitrazepam | C16H14FN3O | 283.29 | 35.47 | 1 | 6 | A6 |
| Cinolazepam | C18H13ClFN3O2 | 243.00 | 30.42 | 1 | 34 | D1 |
| Flurazepam | C21H23ClFN3O | 233.55 | 29.24 | 1 | 22 | B11 |
| Bromazepam | C14H10BrN3O | 200.69 | 25.13 | 1 | 7 | A7 |
| Diclazepam | C16H12Cl2N2O | 191.88 | 24.02 | 1 | 12 | B1 |
| Halazepam | C17H12ClF3N2O | 170.83 | 21.39 | 1 | 28 | C6 |
| Cloniprazepam | C19H16ClN3O3 | -36.71 | -4.60 | 1 | 40 | D7 |
| Prazepam | C19H17ClN2O | -48.00 | -6.01 | 1 | 31 | C9 |
| Flutoprazepam | C19H16ClFN2O | -142.26 | -17.81 | 1 | 45 | E1 |
| Flunitrazolam | C17H12FN5O2 | 1094.57 | 137.03 | 2 | 41 | D8 |
| Fluclotizolam | C15H10ClFN4S | 633.93 | 79.36 | 2 | 43 | D10 |
| Clonazolam | C17H12ClN5O2 | 609.21 | 76.27 | 2 | 26 | C4 |
| Flubromazolam | C17H12BrFN4 | 466.74 | 58.43 | 2 | 25 | C3 |
| Nitrazolam | C17H13N5O2 | 450.86 | 56.44 | 2 | 37 | D4 |
| Flualprazolam | C17H12ClFN4 | 444.31 | 55.62 | 2 | 48 | E4 |
| Estazolam | C16H11ClN4 | 408.62 | 51.16 | 2 | 15 | B4 |
| 1-demethyl Phenazolam | C16H10BrClN4 | 327.31 | 40.98 | 2 | 44 | D11 |
| Phenazolam | C17H12BrClN4 | 305.76 | 38.28 | 2 | 49 | E5 |
| Midazolam | C18H13ClFN3 | 285.67 | 35.76 | 2 | 23 | C1 |
| Triazolam | C17H12Cl2N4 | 281.71 | 35.27 | 2 | 36 | D3 |
| Alprazolam | C17H13ClN4 | 255.21 | 31.95 | 2 | 5 | A5 |
| Bromazolam | C17H13BrN4 | 252.71 | 31.64 | 2 | 38 | D5 |
| Etizolam | C17H15ClN4S | 243.64 | 30.50 | 2 | 4 | A4 |
| 1'-hydroxy Midazolam | C18H13ClFN3O | 242.79 | 30.40 | 2 | 1 | A1 |
| 4-hydroxy Alprazolam | C17H13ClN4O | 234.12 | 29.31 | 2 | 13 | B2 |
| Pyrazolam | C16H12BrN5 | 228.95 | 28.66 | 2 | 9 | A9 |
| α-hydroxy Alprazolam | C17H13ClN4O | 228.00 | 28.54 | 2 | 11 | A11 |
| 7-Aminoclonazolam | C17H14ClN5 | 208.88 | 26.15 | 2 | 50 | E6 |
| Adinazolam | C19H18ClN5 | 207.05 | 25.92 | 2 | 27 | C5 |
| Ketazolam | C20H17ClN2O3 | -7.98 | -1.00 | 2 | 2 | A2 |
| Icilin (100uM) | | 798.76 | 100.00 | - | 55 | E11 |

**Supplementary Figure 1**

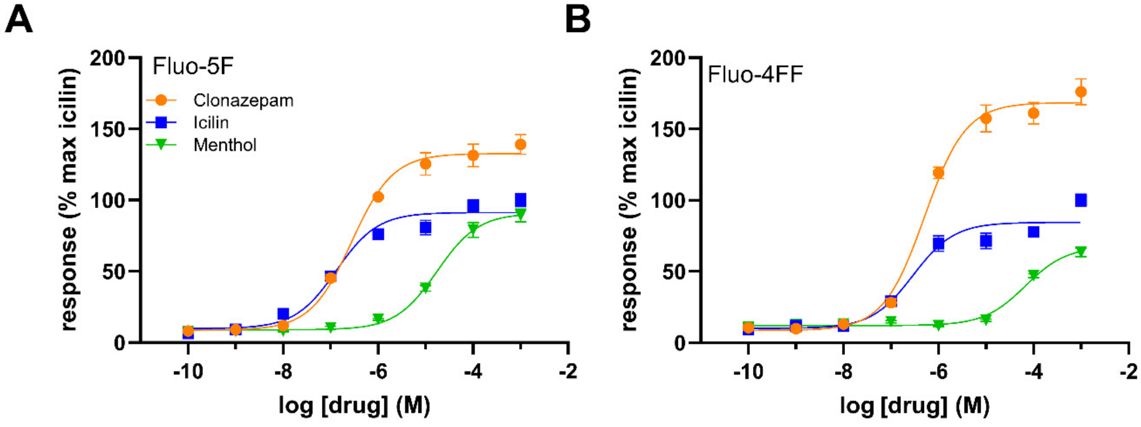


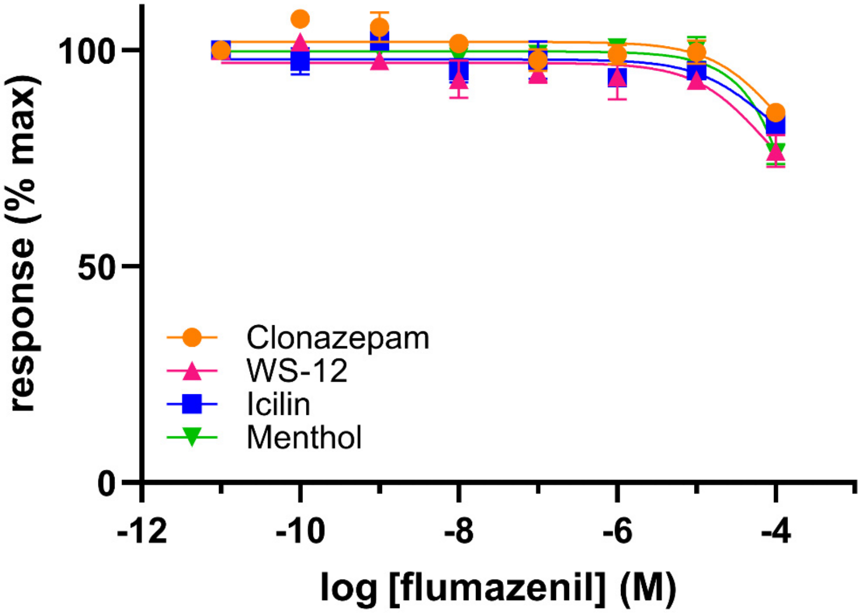

**Supplementary Figure 2**

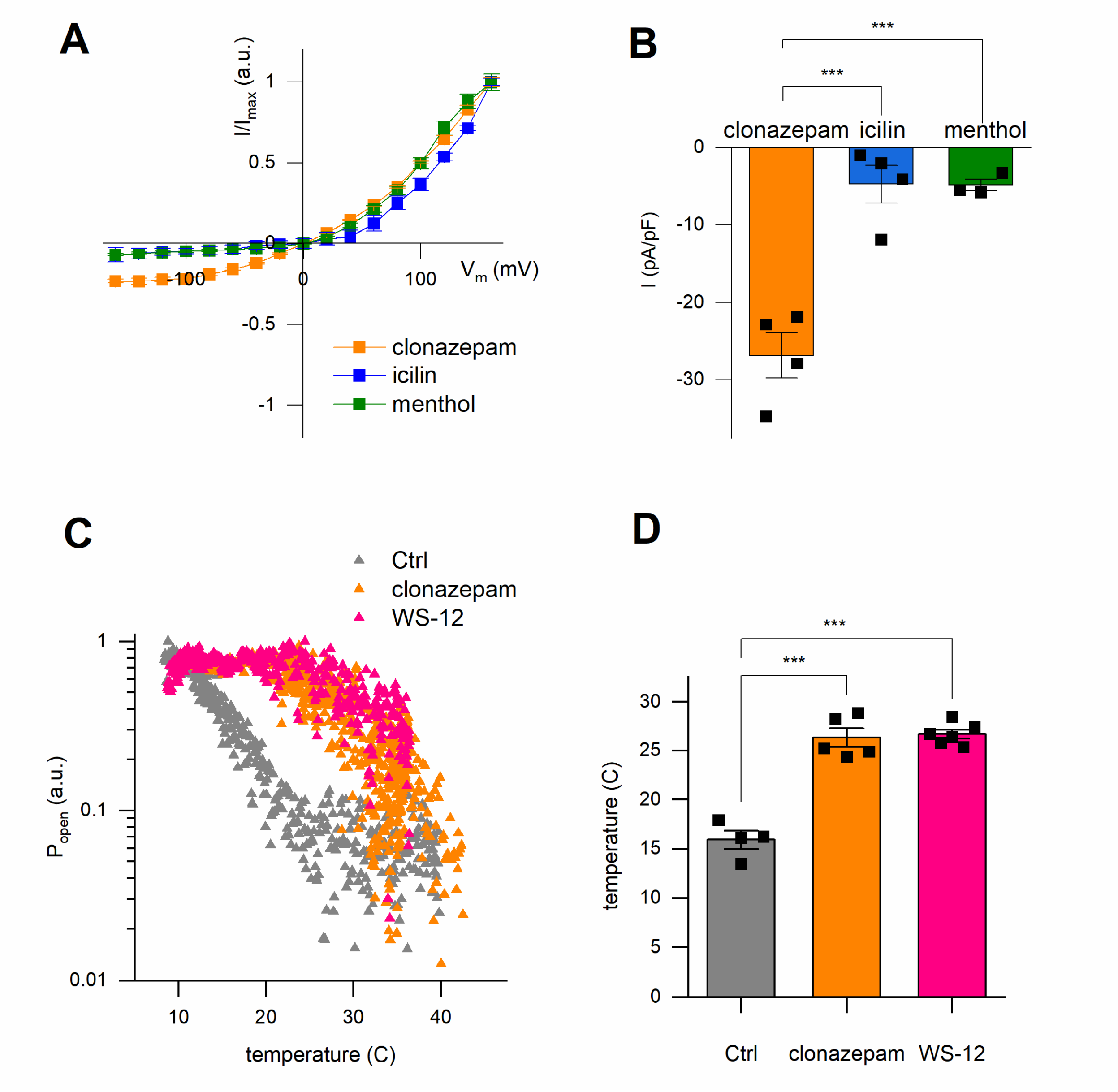

**Supplementary Figure 3**

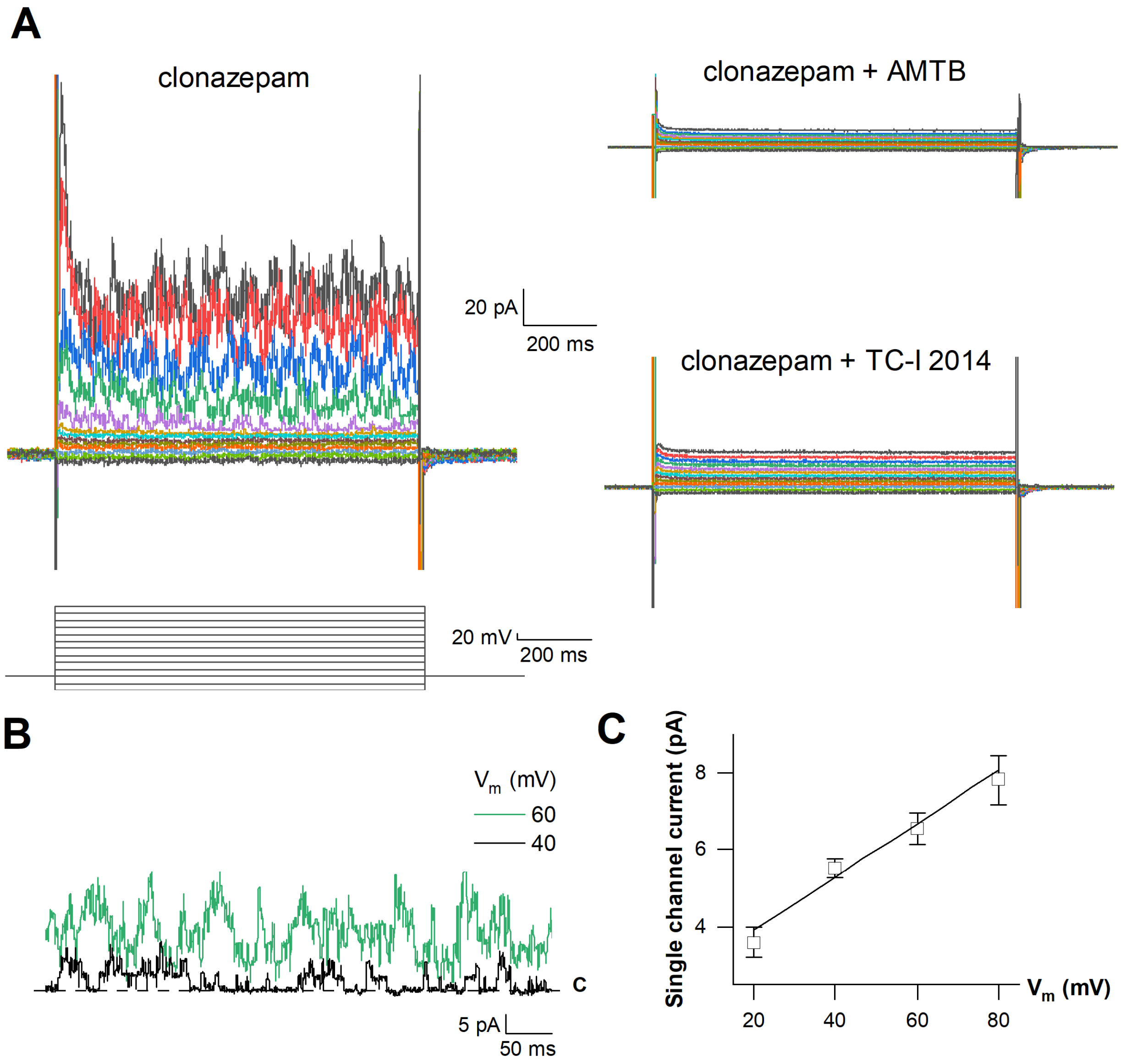

**Supplementary Figure 4**

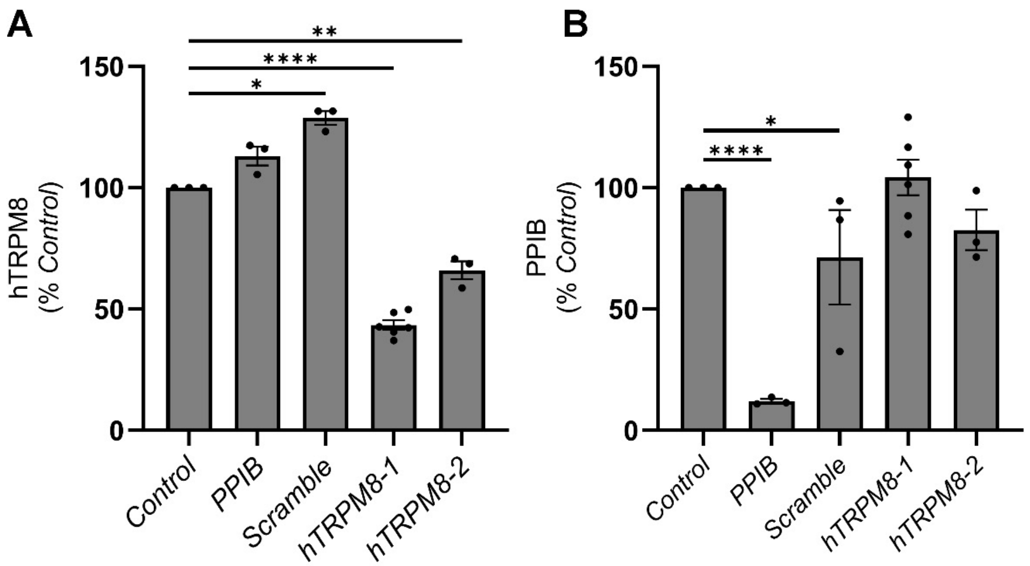
